# Information storage across a microbial community using universal RNA memory

**DOI:** 10.1101/2023.04.16.536800

**Authors:** Prashant B. Kalvapalle, August Staubus, Matthew J. Dysart, Lauren Gambill, Kiara Reyes Gamas, Li Chieh Lu, Jonathan J. Silberg, Lauren B. Stadler, James Chappell

## Abstract

Biological recorders can code information in DNA, but they remain challenging to apply in complex microbial communities. To program microbiome information storage, a synthetic catalytic RNA (cat-RNA) was used to write information in ribosomal RNA (rRNA) about gene transfer host range. By reading out native and modified rRNA using amplicon sequencing, we find that 140 out of 279 wastewater microbial community members from twenty taxonomic orders participate in conjugation and observe differences in information storage across amplicon sequence variants. Twenty of the variants were only observed in modified rRNA amplicons, illustrating information storage sensitivity. This autonomous and reversible RNA-addressable memory (RAM) will enable biosurveillance and microbiome engineering across diverse ecological settings and studies of environmental controls on gene transfer and cellular uptake of extracellular materials.

**One-Sentence Summary:** Ribosomal RNA sequencing detects cellular events recorded across a wastewater microbial community using synthetic biology.

## MAIN TEXT

Nucleic acids acquired through gene transfer can alter cell-cell interactions (*1, 2*), facilitate niche expansion (*3*), drive bacterial resistance to phage (*4*), and control gene stability (*5*). Gene transfer is also critical to microbial domestication for synthetic biology (*6*–*8*) and a challenge for the safe application of such technologies (*9*). Because gene transfer can occur across species from different genera, phyla, and even kingdoms (*10, 11*), there is a need to understand how it varies across communities. Of particular importance is understanding the host ranges of conjugative plasmids (*12*), bacteriophages (*13*), and environmental DNA uptake (*10*), as well as the effects of biotic and abiotic parameters on gene transfer rates (*14*–*19*). Such knowledge is crucial to prevent the unwanted spread of harmful antibiotic resistance genes (*20*) and DNA from genetically-engineered biotechnologies (*21*).

Genetically-encoded reporters are commonly used to study gene transfer (*22, 23*). Fluorescent proteins and antibiotic resistance genes can be coded into mobilizable genetic elements and used to identify cells that participate in gene transfer (*24*). These approaches led to the discovery of novel DNA exchange mechanisms (*25, 26*) and provided mechanistic insight into gene transfer processes (*27*). Since these methods require microbial growth and propagation, or conditions amenable to optical outputs for fluorescent reporters, they cannot be applied in native communities growing within non-transparent environmental materials, where most microbes reside on Earth (*28*), and which influence gene transfer rates (*29*).

Culture-independent strategies can autonomously record information about gene transfer in a community without the need for imaging. For instance, mobile DNA containing a transposase and transposon can be used to randomly integrate a transposon into the genomes of cells that take up genetic material (*30*). However, this approach requires metagenomic sequencing to detect the transposon and identify transconjugants, limiting sensitivity, it cannot measure plasmid dynamics as the insertion is irreversible, and it depends upon transposase transcription, translation, and folding (*31*), which varies across microbes. Enzymatic strategies for genetic barcoding overcome some of these limitations (*32*), but they cannot be performed autonomously within living cells and require arduous chemical manipulations.

To monitor gene transfer across an environmental microbiome, we designed catalytic RNAs (cat-RNA) that autonomously barcode host RNA upon gene transfer without the need for translation (Fig. 1A). These small, genetically-encoded cat-RNA are composed of: (i) a designable *<u>RNA guide</u>* that localizes the system to a target RNA through base pairing interactions, (ii) a *<u>catalytic core</u>* derived from a ribozyme that serves as a writer to catalyze the barcoding reaction (*33, 34*), and (iii) a non-coding *<u>RNA barcode</u>*.

**Figure 1.**
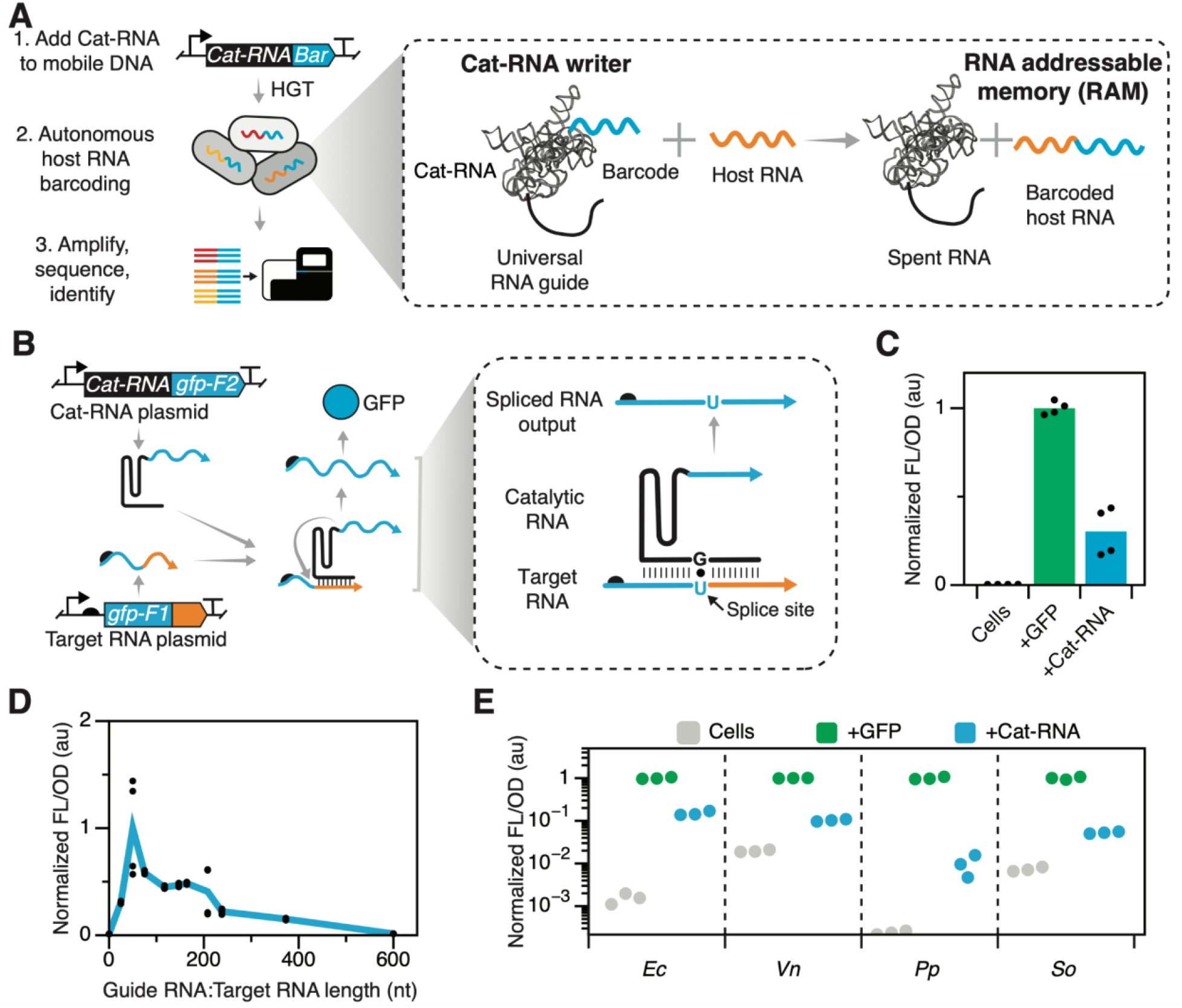
RNA-Addressable Memory (RAM) is an efficient barcoding technology that functions across diverse microbes. (**A**) RAM uses a cat-RNA to splice synthetic RNA barcodes onto host RNA to record mobile DNA uptake by horizontal gene transfer (HGT). (**B**) A fluorescence assay was created that uses cat-RNA barcoding of a target RNA to produce a mRNA product that is competent for translating GFP. (**C**) Fluorescence of *E. coli* expressing the cat-RNA reporter (+Cat-RNA) or a constitutively-expressed GFP positive control (+GFP) is compared with cells harboring an empty vector (Cells). The signal from cat-RNA is significantly higher than the vector control (two-tailed, unpaired t test, p<0.05). (**D**) RNA guide length affects the reporter signal. RNA guides and corresponding target RNA, ranging from 25 to 373 nt, present signals that are significantly higher than a cat-RNA design lacking a guide (two-tailed, unpaired t test, p<0.05). (**E**) Fluorescence characterization of the visual reporter in *E. coli* (*Ec*), *V. natriegens* (*Vn*), *P. putida* (*Pp*), and *S. oneidensis* (*So*). All cells present fluorescence (+Cat-RNA) that is significantly higher than the negative control (Cells), like the positive control (+GFP) (two-tailed, unpaired t test, p<0.05). Three or more biological replicates were acquired for each experiment.

Upon interaction with the target RNA, cat-RNAs are designed to amend the RNA barcode onto the target RNA, forming a genetically-encoded memory (Supplementary Fig. 1), which we designate RNA-addressable memory (RAM).

To determine cat-RNA efficiencies across different microbes, we developed a split *gfp* gene that functions as a visual reporter for RNA splicing (Fig. 1B). The first *gfp* fragment, which is translated, contains the coding sequence for GFP residues 1 to 65 (gfp-F1) fused to a uracil that is targeted for splicing and followed by a non-coding RNA (ncRNA) sequence. The second *gfp* fragment, which is fused to the end of a cat-RNA writer, has a guide that targets the last six nucleotides (nt) of gfp-F1 and 50 nt of the ncRNA sequence. The latter cat-RNA writer amends an RNA barcode composed of the coding sequence for GFP residues 66-238 (gfp-F2) to the end of gfp-F1 to create a native GFP transcript. When these RNA were transcribed in *Escherichia coli*, whole cell fluorescence exceeded that of cells transformed with empty vector (cells) and was ∼30% of that in cells that constitutively express native GFP (Fig. 1C, Supplementary Fig. 2). The growth rates of cells expressing the reporter were not significantly different from cells lacking it (Supplementary Fig. 3). These results show that cat-RNA efficiently generates chimeric transcripts through RNA-mediated barcoding without imposing a cellular burden.

**Figure 2.**
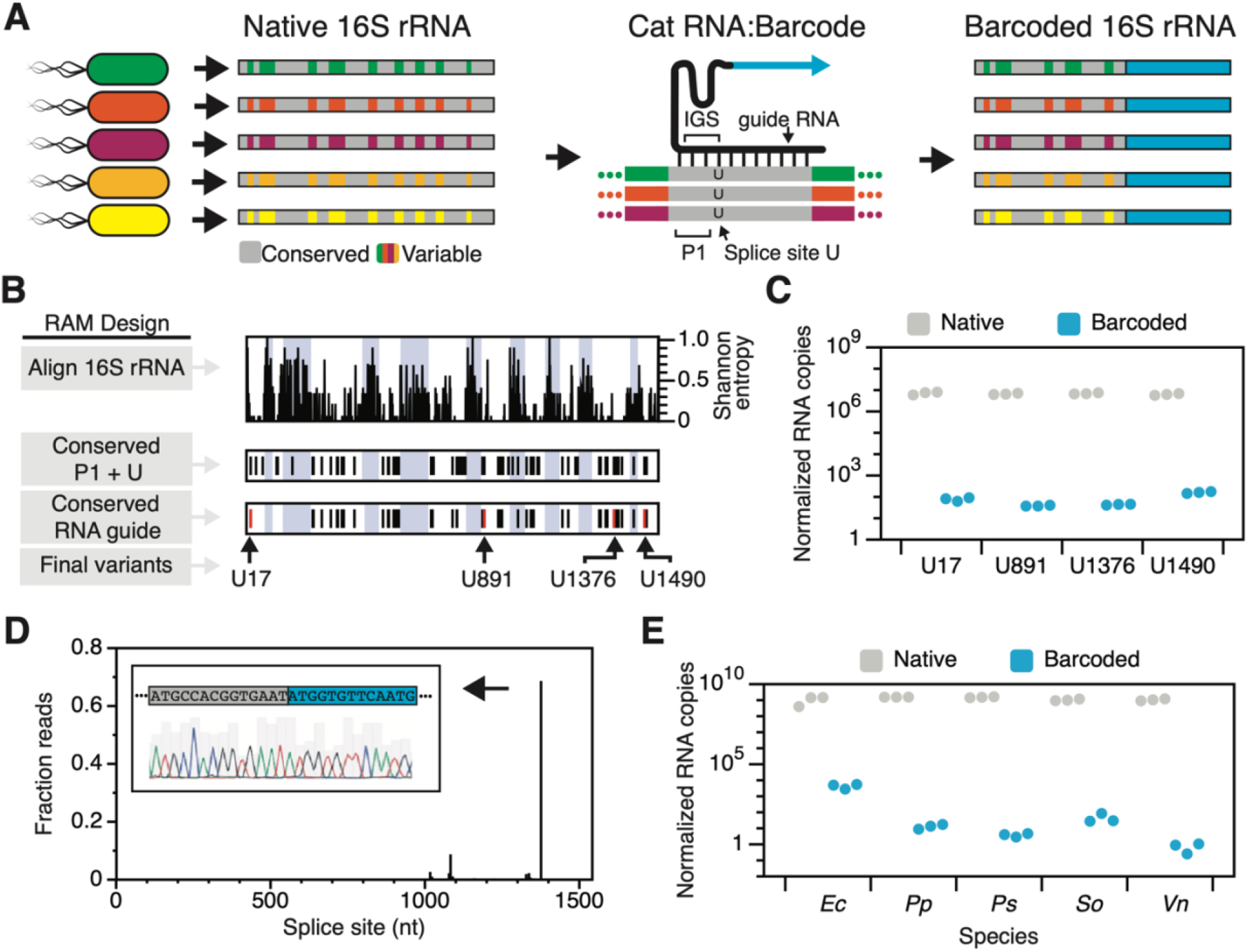
Universal cat-RNA can record information in conserved 16S rRNA sequences. (**A**) Cat-RNAs that target conserved regions of 16S rRNA (gray) add barcodes (blue) such that amplicon sequencing can identify the cells that participate in gene transfer (variable). In 16S rRNA, P1 sequences adjacent to the targeted uracils (U) form duplex interactions with the internal guide sequence (IGS) in cat-RNAs. (**B**) Bioinformatic analysis of 16S rRNAs reveal the Shannon entropy, potential P1 and U splice sites conserved across microbes, and guide RNA targets having high annealing strengths. The designs tested are in red. (**C**) Quantification of native and barcoded 16S rRNA in *E. coli* expressing each cat-RNA using RT-qPCR. (**D**) The cat-RNA designed to target U1376 generates the expected barcoded product, revealed by amplicon and Sanger sequencing (inset), with minimal non-specific barcoding. The sequence shown is from a single amplicon. (**E**) Quantification by RT-qPCR of native and barcoded 16S rRNA in *E. coli* (*Ec*), *P. putida* (*Pp*), *P. stutzeri* (*Ps*), *S. oneidensis* (*So*), and *V. natriegens* (*Vn*) transformed with plasmids encoding the cat-RNA that targets U1376. Data represent three biological replicates.

**Figure 3.**
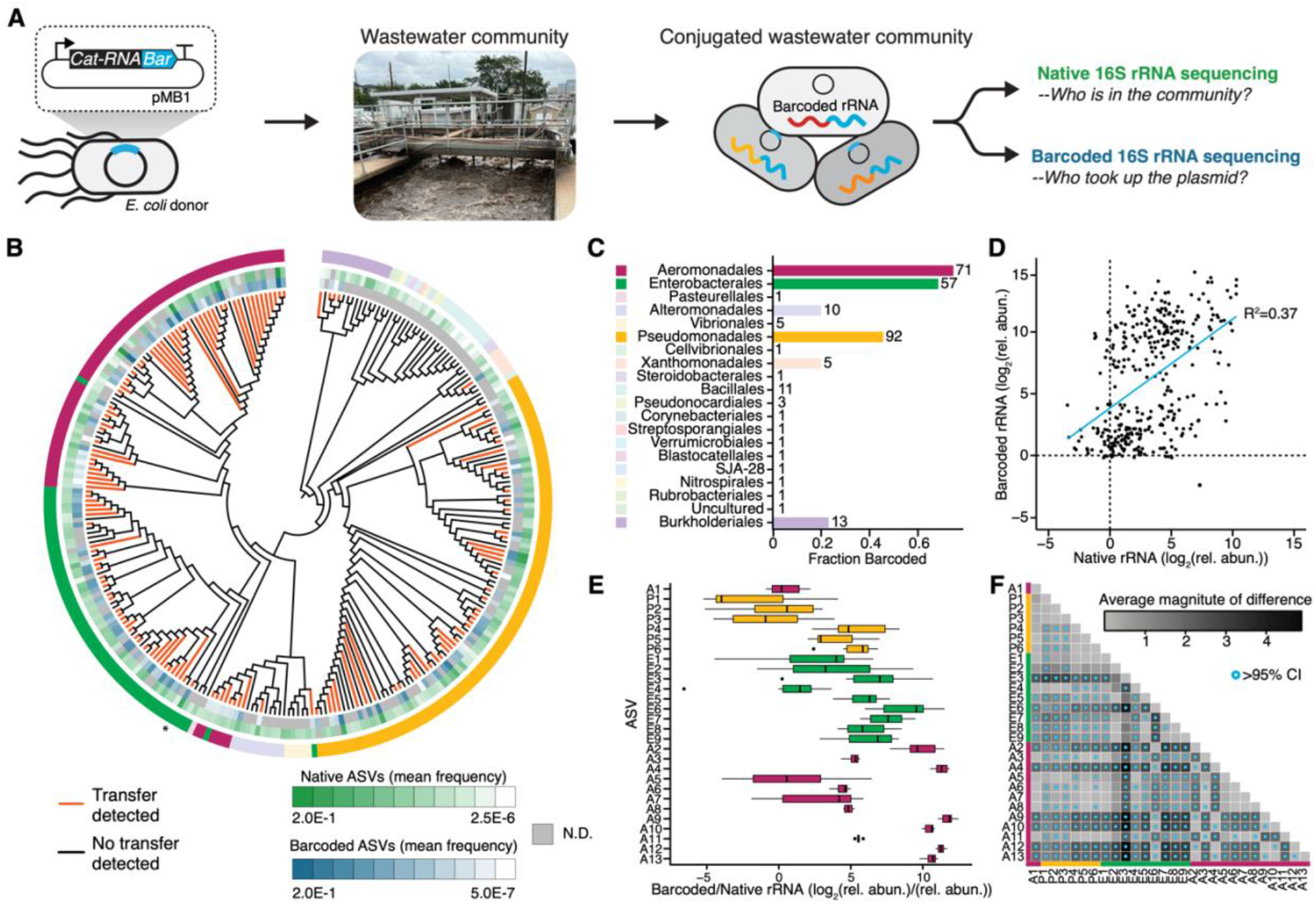
Information storage in a Houston wastewater microbial community. (**A**) RAM was used to record information about conjugation within a Houston wastewater microbial community, and amplicon sequencing of the native and barcoded 16S rRNA showed which microbes were present and who participated in conjugation. (**B**) Conjugation host range (orange) is mapped onto an evolutionary tree including all ASVs observed in the community. The relative abundances of native (green) and barcoded (blue) ASVs are shown, with the outer leaves showing taxonomic order. The asterisk indicates the position of the *E. coli* donor on the tree. (**C**) For each order, the fraction of total ASVs that participated in conjugation is shown, as well as the number of ASVs. (**D**) A comparison of native and barcoded 16S rRNA frequencies from individual biological replicates reveals a weak linear correlation (y = 3.5 + 0.9x; R^2^ = 0.37). (**E**) For those ASVs that yielded detectable native and barcoded 16S rRNA in all six biological replicates, the median ratio of barcoded/total 16S rRNA is shown using whisker plots. (**F**) Pairwise differences in the barcoded/native 16S rRNA ratios. Bootstrapping was used to determine which differences lie outside of the 95% confidence intervals (CI) (blue points).

We next investigated how RNA guide length affects barcoding. Cat-RNA writers that generate a GFP output were designed with RNA guides ranging in length from 0 to 600 nt. A plasmid expressing each cat-RNA and target RNA was transformed into *E. coli*, and cellular fluorescence was measured (Fig. 1D). A wide range of guide lengths (25 to 375 nt) presented a signal, with the highest occurring with the 50 nt guide. To determine if the cat-RNA writers function in different microbes, we introduced a broad host plasmid encoding the cat-RNA reporter into microbes from diverse environments, including gut (*E. coli*), ocean (*Vibrio natriegens*), soil (*Pseudomonas putida*), and freshwater (*Shewanella oneidensis* MR1). In all cases, a fluorescence signal was observed for the cat-RNA reporter (Fig. 1E). Thus, a single cat-RNA writer functions across microbes from different orders without optimization of the cat-RNA sequence or the transcription cassette.

Given the centrality of 16S rRNA in taxonomic analysis, we hypothesized that a universal cat-RNA writer could be created by modifying the RNA guide so that it is complementary to conserved 16S rRNA regions (Fig. 2A). To design universal cat-RNAs, we performed a 16S rRNA multiple sequence alignment using Gram negative and positive microbes to identify conserved regions (6 nt) that end in a uracil (Fig. 2B). The 5 nt adjacent to the uracil will ultimately bind to the internal guide sequence (IGS) to form a paired domain called P1 (*35*–*37*). We then designed RNA guide sequences that are complementary to the rRNA sequences adjacent to each conserved IGS-U (Supplementary Fig. 4). We scored each variant based on conservation of the guide-binding region and identified 54 designs across the targeted species with >80% identity (Supplementary Fig. 5). This finding shows a universal RNA guide can be designed to target cat-RNA to diverse 16S rRNA sequences for information storage.

Four universal cat-RNA writers were built that target conserved rRNA sequences (Supplementary Fig. 6). These cat-RNAs were expressed from plasmids in *E. coli* using a constitutive promoter (Fig. 2C), and the native and barcoded 16S rRNA was quantified by reverse transcription-quantitative polymerase chain reaction (RT-qPCR). All four of the designs produced barcoded 16S rRNA. A comparison of the barcoded 16S rRNA to total rRNA abundance revealed that these cat-RNAs presented similar barcoding efficiencies, with 5 to 28 barcoded-rRNA per million total rRNA molecules. Assuming that exponentially growing cells contain ∼72,000 copies of rRNA (*38*), this finding shows that 0.4 to 2 copies of rRNA per cell are barcoded.

To establish the sequence of the barcoded rRNA, we performed RT-PCR and amplicon sequencing of the spliced product arising from the cat-RNA that targets U1376 (Fig. 2D). The major product arose from ligation of the barcode after U1376, although low frequency off-pathway splicing occurred at sites with uracils and related P1 sequences (Supplementary Fig. 7). We next investigated the stability and the reversibility of the barcoded rRNA (Supplementary Fig. 8). This analysis revealed stability that is similar to that of a typical bacterial mRNA (*39*). Taken together, these findings show that cat-RNA can barcode a conserved rRNA sequence and that information storage represents a reversible form of biological memory. The cat-RNA that targets U1376 was used for all subsequent analyses.

To determine whether cat-RNA efficiently barcodes rRNA in different microbes, we cloned it onto a broad host plasmid (pBBR1) and transformed this plasmid into *E. coli, P. putida, Pseudomonas stutzeri, S. oneidensis*, and *V. natriegens*. RT-qPCR detected barcoded rRNA was detected in all five species although species-to-species variation in the fraction of barcoded 16S rRNA was observed (Fig. 2E). This variation correlated (R^2^> 0.99) with cat-RNA transcription in each microbe (Supplementary Fig. 9). These findings demonstrate that a universal cat-RNA coded into a single broad-host-range plasmid can record information within a conserved rRNA sequence in diverse microbes, and they show how the cat-RNA signal can be used to rapidly compare transcription from the same promoter across different organisms.

The finding that cat-RNA can barcode rRNA in a range of microbes suggested it could be applied in a microbiome from the environment to record information about who participates in gene transfer. Because 16S rRNA sequences are used for taxonomic classification, we hypothesized that cat-RNA would generate an ensemble of chimeric rRNA that could be sequenced and analyzed to identify the organisms that participated in gene transfer. Since conjugation rates can vary widely (*40*), we programmed *E. coli* to minimize the donor rRNA barcoding signal by repressing cat-RNA transcription >10^3^ fold (Supplementary Fig. 10). We used this strain as a donor for conjugation into a wastewater sludge microbial community sampled from the West University Wastewater Treatment Plant in Houston, Texas (Fig. 3A). Filter mating assays were performed aerobically, total RNA was isolated and converted to cDNA, and amplicon sequencing was performed. Analysis of barcoded and native rRNA revealed 279 amplicon sequence variants (ASVs), with microbes from twenty orders (Fig. 3B). Among all ASVs detected, 140 presented 16S rRNA-barcode signals. Among these ASVs, twenty were only detected when sequencing the barcoded rRNA (Supplementary Figure 11). Thus, amplicon sequencing can be used with cat-RNA to read out information about who is present in a community and who participates in gene transfer.

To understand how ASV relatedness correlates with information storage, we quantified the fraction of ASVs barcoded across each order (Fig. 3C). A majority of the Proteobacteria (60%), which represented almost 90% of the ASVs in the consortium, contained barcoded 16S rRNA. The order *Aeromonadales* presented the largest fraction of barcoded ASVs (∼70%). The barcoding signal decreased with ASV taxonomic distance from this order, as did ASV abundance in each order. This data shows that diverse microbes in the wastewater consortia are capable of information storage about DNA uptake and participation in gene transfer, and it shows that *Aeromonadales, Enterobacterales*, and *Pseudomonadales* are active participants in gene transfer, consistent with prior studies (*3, 41, 42*).

To evaluate whether ASV abundance affects barcoding, we analyzed the relationship between total and barcoded 16S rRNA for the ASVs that presented detectable levels of both (Fig. 3D). This analysis yielded a linear trend (m = 0.9, R^2^ = 0.37), suggesting that microbial abundance in a consortium contributes to the barcode signal. However, the slope deviates from unity, suggesting ASV abundance is not the only control on the signal. Some ASVs presented a barcoded rRNA signal, even though their native 16S rRNA was not detected (Supplementary Figure 12). This latter observation shows the exquisite sensitivity of RNA memory to store and read out information about low abundance consortium members participating in gene transfer.

A comparison of wastewater microbiome samples containing or lacking a non-native microbe revealed the variance of our experiment (Supplementary Figure 13). To investigate if we could detect variation in barcoding efficiency across ASV pairs, we compared the ratios of the barcoded to total 16S rRNA amplicon frequencies. When this analysis was performed across only those ASVs that had detectable barcoded and total rRNA in all six replicates (Fig. 3E), we found that the ratio varied by up to ∼43,000 fold. A pairwise analysis of this data revealed a range of differences in their ratios across ASVs within the same order and between different orders (Fig. 3F). Bootstrap analysis of the data revealed that many pairwise comparisons presented differences that were outside of the 95% confidence intervals (Supplementary Figure 14). These findings show that closely related organisms present variation in barcoding, which is thought to arise from differences in ASV conjugation rates, promoter activities, and the stabilities of cat-RNA and the mobile element coding for cat-RNA.

The facile information storage enabled by cat-RNA has several advantages compared with existing memory systems (*43*–*49*). First, cat-RNA records information within universally-expressed biomolecules at highly conserved sequences adjacent to variable sequences that can be used to distinguish taxa (*50*). As such, well-established rRNA sequencing pipelines used for environmental microbiology can be used to read out consortium information (*51*). Second, cat-RNA uses a small biomolecular writer (<500 nt) that only requires a single promoter to synthesize, while existing autonomous barcoding systems require transcription and translation (*30*). Thus, RNA memory can easily store information in microbes across a complex microbiome by coupling cat-RNA synthesis to broad-host-range promoters (*52*). Third, the use of RNA for writing and storing information is expected to minimize resource burdens, since the cost of protein synthesis required by existing memory systems is much greater than RNA synthesis (*53*). Fourth, after cells are cured of a mobile genetic element expressing cat-RNA, the recorded information will not be vertically inherited, making it compatible with dynamic information storage. This can be contrasted with DNA memory devices, such as lineage tracers (*54*), whose power lies in the vertical inheritance of information.

To diversify this proof-of-concept RNA memory for information storage, alternative promoters can be used to regulate cat-RNA synthesis, and donor strains that generate different DNA modifications can be applied to minimize mobile DNA degradation by restriction-modification systems (*55*). Alternative RNA guides can be created to selectively store information in a subset of consortia members by targeting RNA sequences that are only conserved in those species. By diversifying RNA barcodes, cat-RNA can be created that are compatible with multiplexing information storage within individual microbes and in consortia. Small, cat-RNA that store information about cellular uptake of molecules across microbial communities will revolutionize our ability to program cells using synthetic biology by allowing researchers to perform the design-build-test-learn cycle in environmental samples, and it will advance our ability to study and understand the environmental controls on the mobility of natural and synthetic DNA that is exchanged via conjugation, transduction, and transformation. Finally, RNA memory should transform our ability to study how environmental conditions affect gene transfer host range and efficiencies across environmental communities.

## ACKNOWLEDGMENTS

We thank Dr. Kreso Josic (University of Houston) for discussions about bootstrap analysis. We thank the West University Wastewater Treatment Plant operators for their assistance with collecting samples.

## AUTHOR CONTRIBUTIONS

Contributions are noted in alphabetical order. Conceptualization: *JC, JJS, LBS*

Methodology: *AS, JC, JJS, KRG, LBS, LCL, LG, MJD, and PBK*

Investigation: *AS, JC, JJS, KRG, LBS, LCL, LG, MJD, and PBK*

Visualization: *AS, JC, JJS, KRG, LBS, LCL, LG, MJD, and PBK*

Funding acquisition: *JC, JJS, and LBS*

Project administration: *JC, JJS, and LBS*

Supervision: *JC, JJS, and LBS*

Writing – original draft: *JC, JJS, and LBS*

Writing – review & editing: *AS, JC, JJS, KRG, LBS, LCL, LG, MJD, and PBK*

## DATA AND MATERIALS AVAILABILITY

Genetic constructs will be made available in Addgene and custom computer scripts uploaded to GitHub. Sequencing data will be made available through NCBI Sequence Read Archive (PRJNA954059).

