## Supplemental information for "Information storage across a microbial community using universal RNA memory"

### TABLE OF CONTENTS


| Item | Title | Page |
|---|---|---|
| Fig. S1 | A comparison of RNA designs with the native ribozyme | 3 |
| Fig. S2 | Single-cell fluorescence analysis of cat-RNA activity | 4 |
| Fig. S3 | Cat-RNA does not affect *E. coli* fitness | 5 |
| Fig. S4 | Sequence conservation in the five nucleotides adjacent to conserved uracils in bacterial 16S rRNAs | 6 |
| Fig. S5 | Comparing different synthetic guide RNA designs | 7 |
| Fig. S6 | Universal cat-RNA guides are mapped onto the structure of 16S rRNA | 8 |
| Fig. S7 | Identification of off-target splicing sites | 9 |
| Fig. S8 | Stability of barcoded rRNA | 10 |
| Fig. S9 | rRNA barcoding correlates with cat-RNA expression | 11 |
| Fig. S10 | Two-plasmid system developed for the conjugative donor cell | 12 |
| Fig. S11 | Evolutionary tree showing the taxonomic rank of wastewater ASVs | 13 |
| Fig. S12 | All ASVs detected from native rRNA and barcoded rRNA sequences | 14 |
| Fig. S13 | Barcoding in the wastewater community containing and lacking a non-native microbe spike in | 15 |
| Fig. S14 | Bootstrapping reveals pairwise differences in ASV barcoding signals | 16 |
| Fig. S15 | Workflow showing data processing to quantify barcoding | 17 |
| Legends | Supplemental Tables 1-4 | 18 |

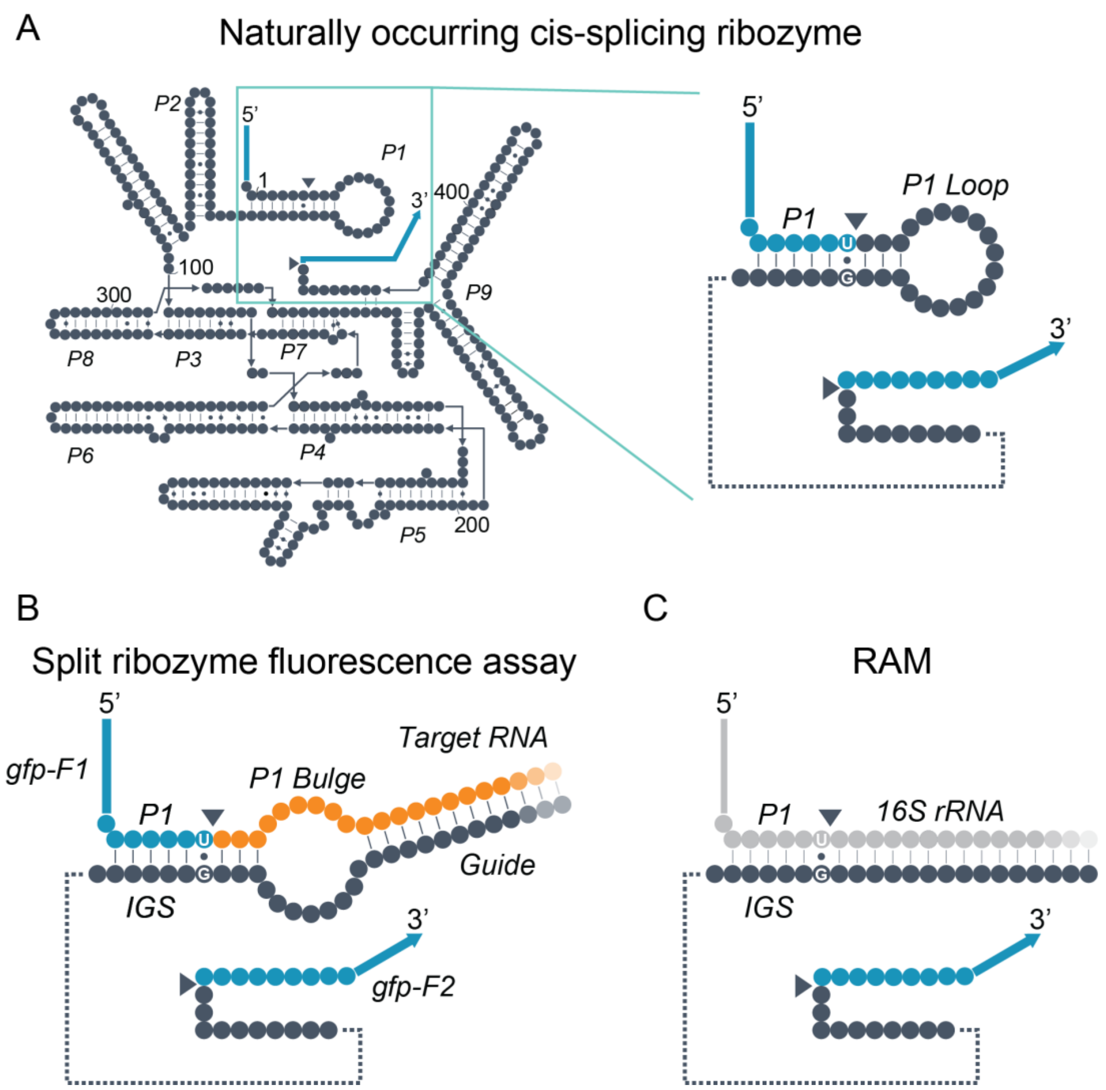


**Supplementary Figure 1. A comparison of RNA designs with the native ribozyme.** (**A**) The native splicing group I intron ribozyme from *Tetrahymena thermophila*. This ribozyme catalyzes a reaction wherein the intron (black) removes itself from the flanking exons (blue, splice sites indicated by triangles) and joins them together into a single RNA strand, which is a *cis*-splicing reaction. (Right panel) the 5' and 3' ends of the intron contain the P1 stem, the P1 loop, and the U•G wobble base pair necessary for splicing. (**B**) Schematic of the synthetic catalytic RNA (cat-RNA) developed as a fluorescence assay for the splicing reaction. (**C**) Schematic of the synthetic cat-RNA that was developed as RNA-addressable memory (RAM), which targets conserved sequences within 16S rRNA for trans splicing reactions.

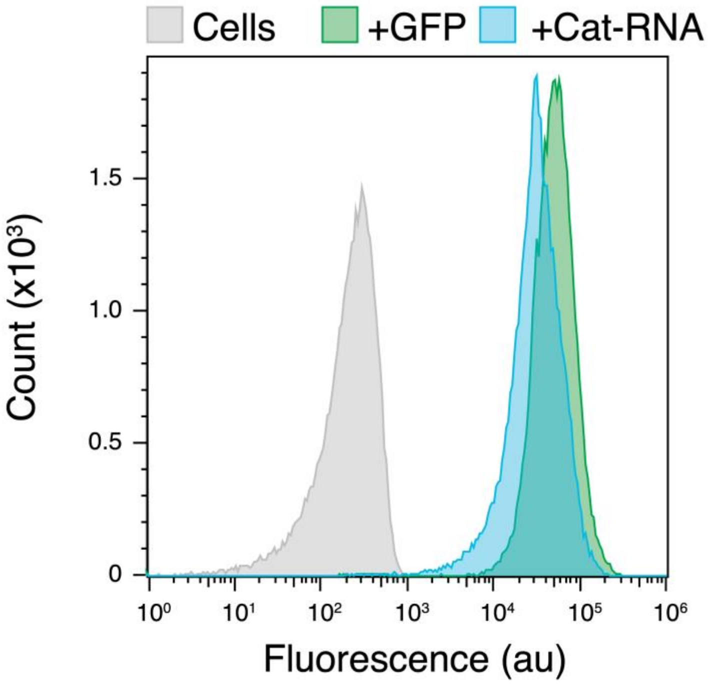


**Supplementary Figure 2. Single-cell fluorescence analysis of cat-RNA activity**. Single-cell GFP fluorescence values of *E. coli* transformed with an empty control plasmid (Cells), a constitutively expressed sfGFP positive control (+GFP), and a plasmid encoding a catalytic RNA (+Cat-RNA) designed to splice two GFP mRNA fragments together as shown in Figure 1B to produce a single mRNA that translates sfGFP. Data were collected from 3 biological replicates. Data shown in units of arbitrary fluorescence (au) are from a single biological replicate that was representative of the other replicates.

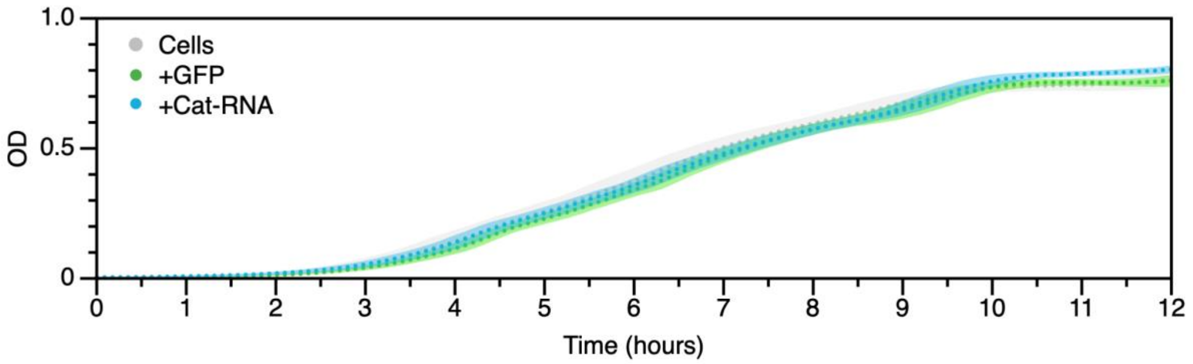


**Supplementary Figure 3. Cat-RNA does not affect *E. coli* fitness**. Optical density (OD) of *E. coli* transformed with an empty control plasmid (Cells), the sfGFP positive control (+GFP), and the plasmid encoding a catalytic RNA (+Cat-RNA) that splices two GFP mRNA fragments together to produce a single mRNA that translates GFP. The maximum specific growth rates ($\mu_{max}$) for cells (0.124 ±0.004 hrs$^{-1}$), GFP (0.125 ±0.003 hrs$^{-1}$), and cat-RNA (0.122 ±0.003 hrs$^{-1}$) presented no significant differences (two-tailed, unpaired *t* test, p>0.05). Data points represent the mean from 12 biological replicates with ±1 s.d. shown as shaded areas.

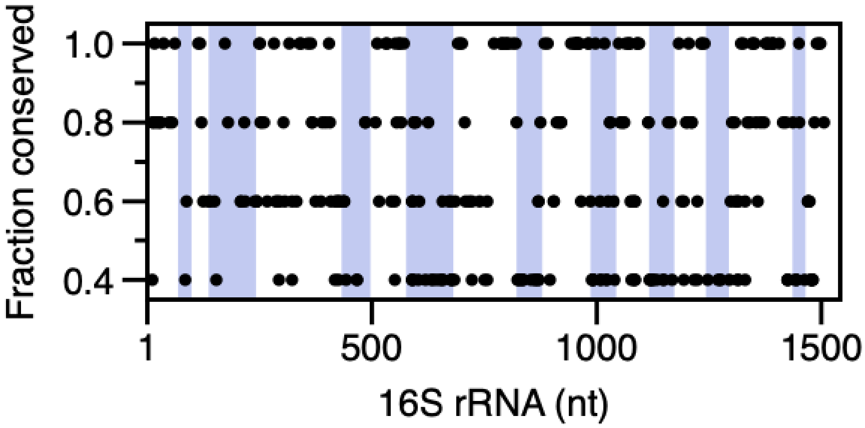


**Supplementary Figure 4. Sequence conservation in the five nucleotides adjacent to conserved uracils in bacterial 16S rRNAs**. To assess the relative quality of different uracils as targets for universal cat-RNA splicing, the 16S rRNA sequences of five microbes (*Escherichia coli*, *Pseudomonas stutzeri*, *Shewanella oneidensis*, *Vibrio natriegens*, and *Bacillus subtilis)* were aligned and the conservation of the five nucleotides (nt) immediately adjacent to conserved uracils found across all species was calculated (Fraction conserved). Only those sequences that were conserved across at least two microbes were considered. In cat-RNA designs (Supplementary Figure 1C), conservation of these nt is critical for forming a P1 stem across all species targeted. Sequence positions are numbered relative to *E. coli* 16S rRNA nt, with the variable regions shaded in blue: V1 (69-99), V2 (137-242), V3 (433-497), V4 (576-682), V5 (822-879), V6 (986-1043), V7 (1117-1173), V8 (1243-1294), and V9 (1435-1465).

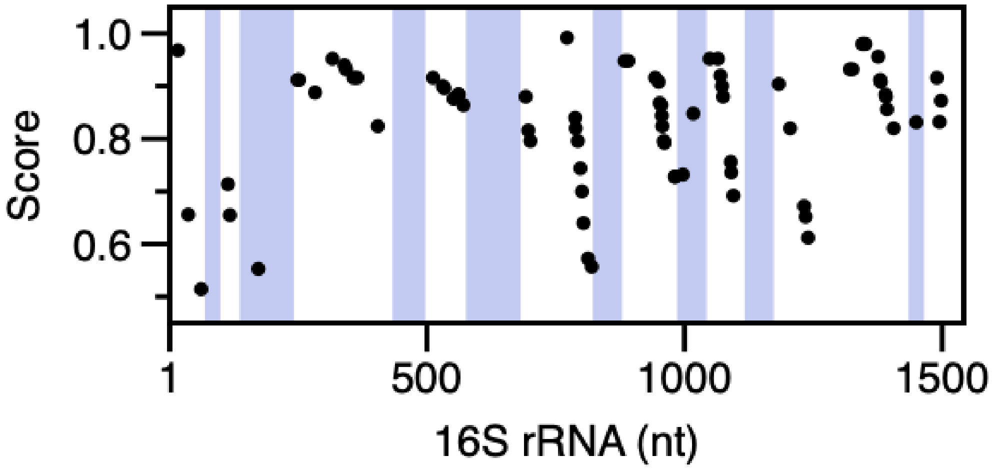


**Supplementary Figure 5. Comparing different synthetic guide RNA designs.** After identifying IGS annealing sites across the five microbes (*E. coli*, *P. stutzeri*, *S. oneidensis*, *V. natriegens*, and *B subtilis)* that are 100% conserved, we assessed whether the 50 nucleotides (nt) downstream of each conserved U and P1 stem represents a good target for a universal cat-RNA guide that targets all five microbes. To do this, we generated a consensus score for the annealing of 50 nt guides to the five different 16S rRNA sequences. Sequence positions are numbered relative to *E. coli* 16S rRNA nt, with the variable regions shaded in blue: V1 (69-99), V2 (137-242), V3 (433-497), V4 (576-682), V5 (822-879), V6 (986-1043), V7 (1117-1173), V8 (1243-1294), and V9 (1435-1465).

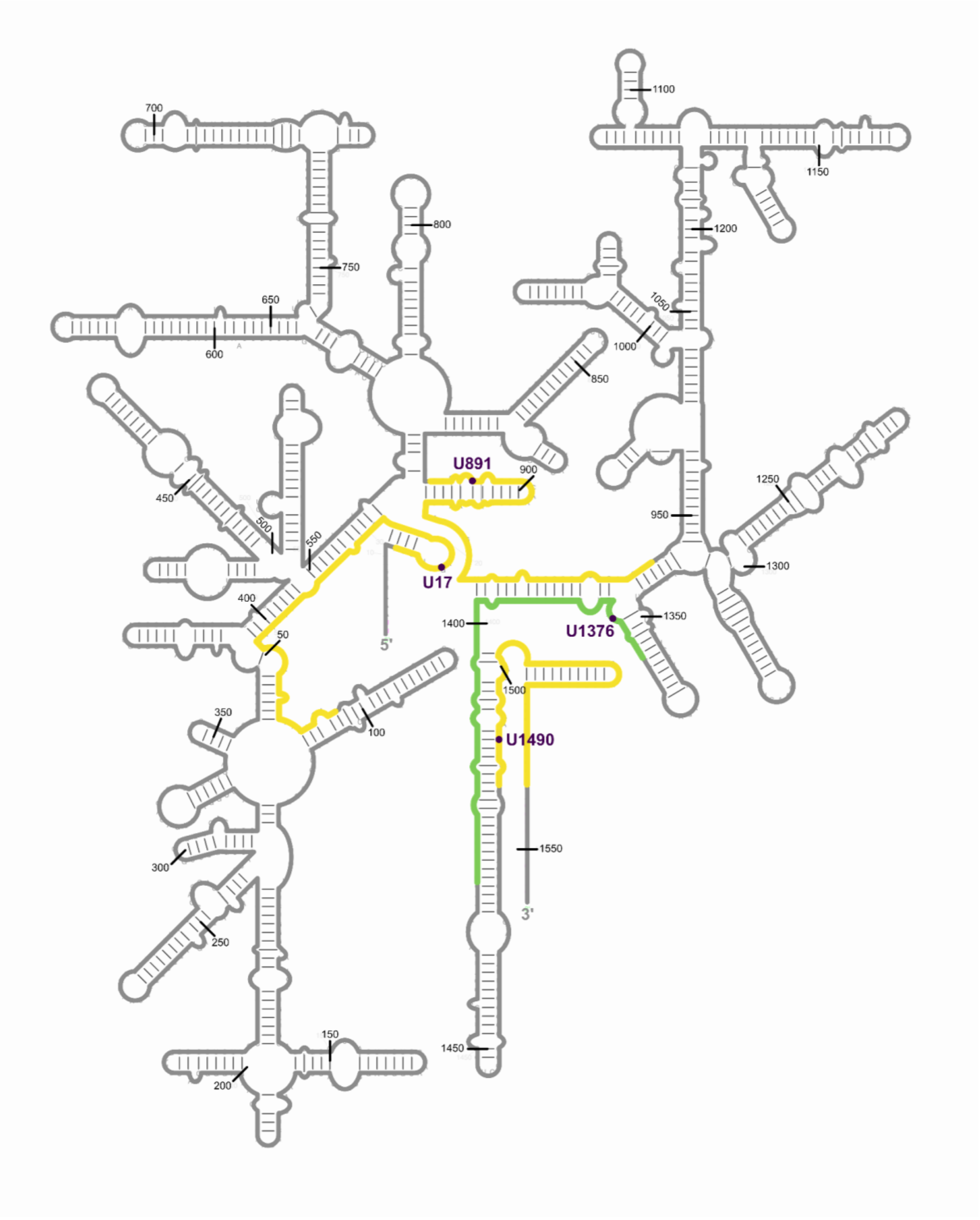


**Supplementary Figure 6. Universal cat-RNA guides are mapped onto the structure of 16S rRNA.** The different cat-RNA guides that were built and tested are mapped onto *E. coli* 16S rRNA. U17, U891, U1376, and U1490 correspond to the splice site uracils targeted by each variant. Yellow indicates regions targeted by the guide sequences in cat-RNA U17, U891, U1376, and U1490. Green indicates the region targeted by cat-RNA U1376, which was used for microbial community barcoding experiments.

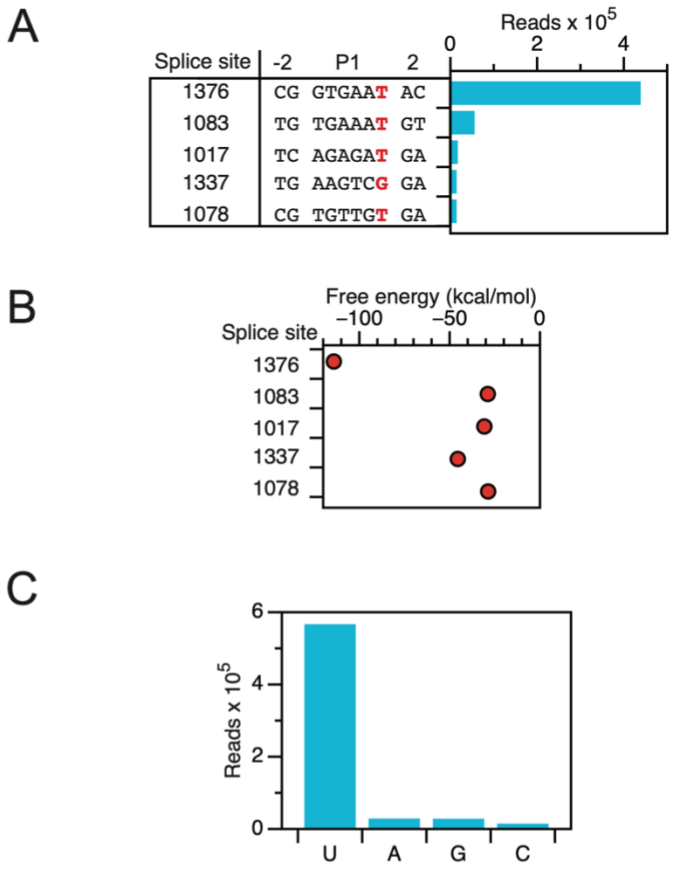


**Supplementary Figure 7. Identification of off-target splicing sites.** (**A**) The relative abundance of the reads observed at each splice site location determined by sequencing barcoded rRNA from *E. coli* expressing cat-RNA U1376. Sequences indicate the 16S rRNA region that forms the P1 stem with the G-IGS (middle) in the cat-RNA, as well as the two nucleotides upstream and downstream of this sequence. The splice site is the 3' nucleotide of the P1 shown (red). The top sequence (splice site 1376) is the expected product. Other products, such as 1017, show some sequence similarity to the canonical P1. (**B**) The predicted free energy of binding between the cat-RNA and the target sequence downstream of each observed splice site shows no correlation between splicing and guide RNA binding energy. Free energy was calculated using the NUPACK algorithm. (**C**) Identity of the observed splice site nucleotide in the 16S rRNA shows the majority of splice sites occur at an uracil, although others can be targeted at low frequencies.

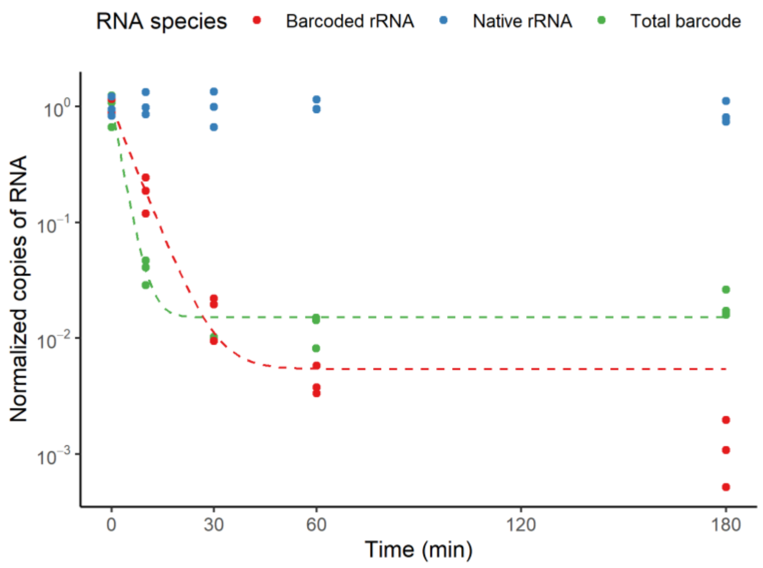


**Supplementary Figure 8. Stability of barcoded rRNA.** The stability of barcoded rRNA was compared with native rRNA and the total barcode using cat-RNA U1376. The total barcode signal can arise from amplification of cat-RNA writer before splicing or barcoded rRNA product (Figure 1A). RT-qPCR was performed on total RNA extracted from *E. coli* containing cat-RNA U1376 at different time points after the addition of rifampicin, which inhibits the initiation of new transcripts by RNA polymerase. Data for each molecular species is normalized to the average RNA copies at the time of rifampicin addition (0 min). Following rifampicin addition, the signal for cat-RNA and barcoded rRNA decreased significantly compared to native rRNA (Welch's one-tailed t-test, $p<0.02$). Data points show the mean and error bars show the s.d. of $n = 3$ independent biological replicates. The curves show exponential fits with $t_{1/2}$ values of 1.9 min (total barcode) and 4 min (barcoded rRNA).

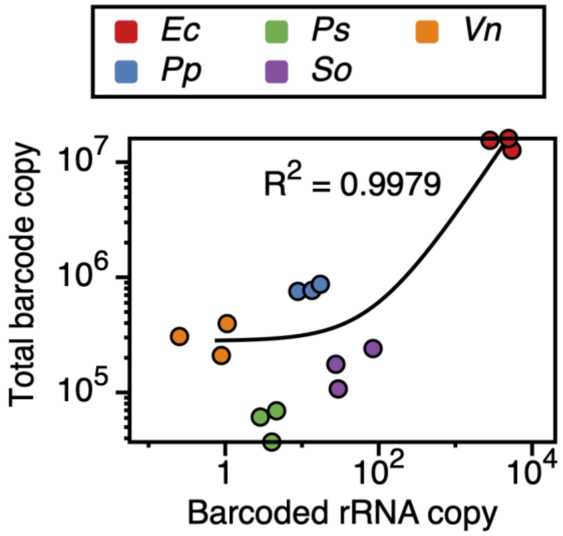


**Supplementary Figure 9. rRNA barcoding correlates with cat-RNA expression.** The total barcode and barcoded rRNA levels were measured by RT-qPCR with standard curve calibration. A comparison of data from five microbes shows a positive correlation between the amount of barcoded rRNA and the transcription level of total barcode. As cat-RNA barcode copy is much greater than barcoded rRNA copy, the total barcode is expected to be correlated to the transcription level of the cat-RNA. A linear fit (y = 280050 + 3280.1*x) is shown. Data represent 3 biological replicates within each organism, including *E. coli* (*Ec*), *P. putida* (*Pp*), *P. stutzeri* (*Ps*), *S. oneidensis* (*So*), and *V. natriegens* (*Vn*).

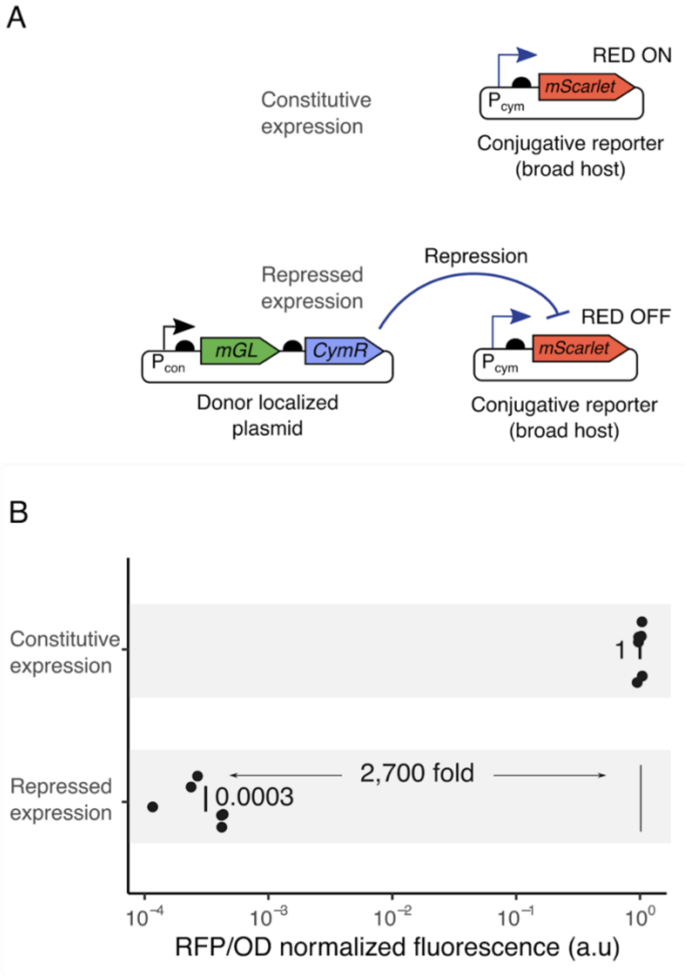


**Supplementary Figure 10. Two-plasmid system developed for the conjugative donor cell.** (**A**) The constructs used to evaluate transcriptional repression of our conjugative plasmid in the donor cells. With this system, the conjugative plasmid is designed to constitutively express mScarlet, while the donor-localized plasmid expresses CymR, which represses mScarlet. (**B**) The relative whole cell fluorescence (RFP/OD) was repressed significantly (Welch's one-tailed t-test, $p = 10^{-8}$), ~2,700 fold in *E. coli* MG1655 harboring the plasmid that expresses CymR. To create a system for conjugation of cat-RNA, the gene encoding mScarlet was replaced with cat-RNA U1376. Data represents the values obtained from six biological replicates, with the means shown as vertical lines.

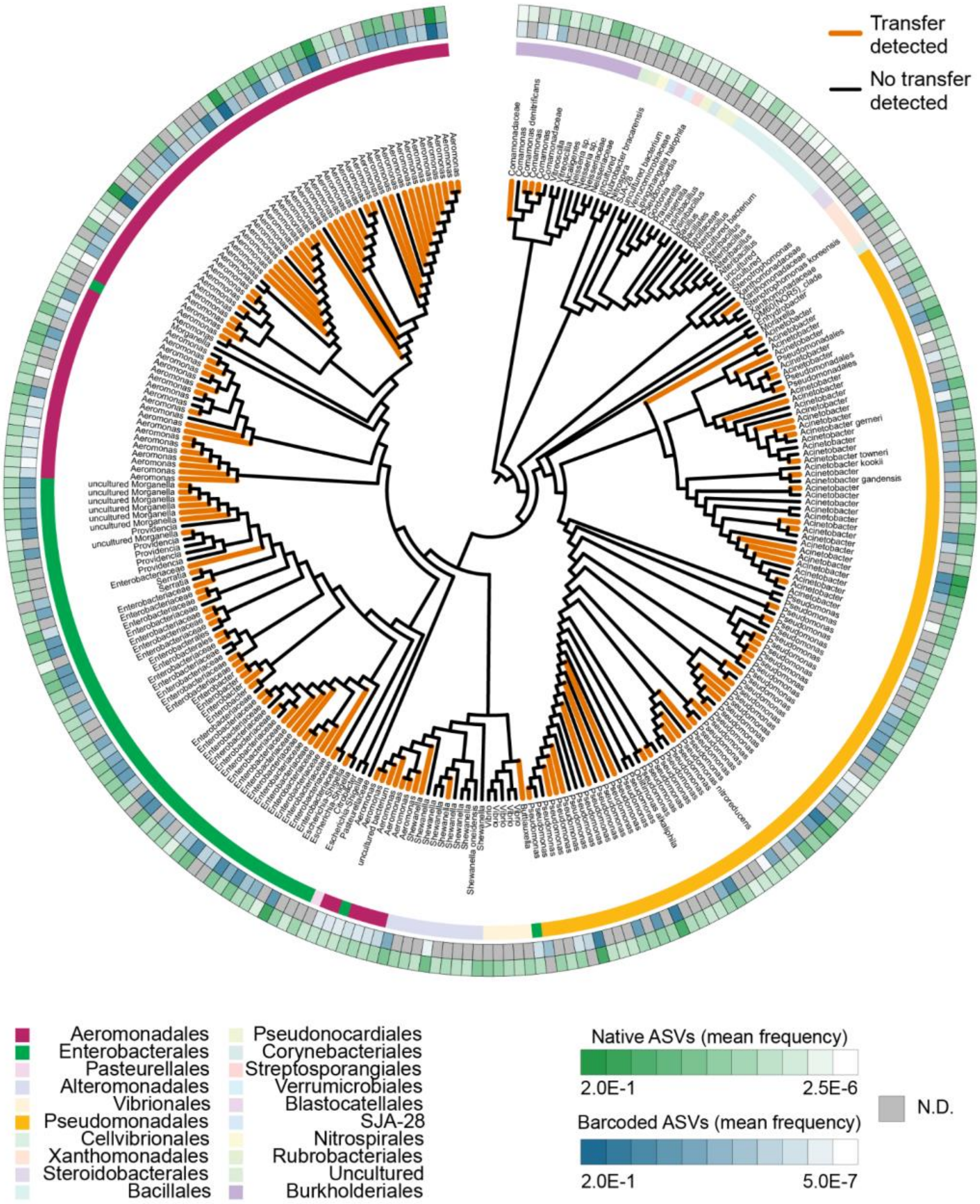


**Supplementary Figure 11. Evolutionary tree showing the taxonomic rank of wastewater ASVs.** The leaves are labeled with the lowest taxonomic rank that could be assigned with ≥70% confidence. Terminal branches are colored orange if the ASV was detected in the barcoded 16S rRNA samples. The innermost circle surrounding the leaves indicates the taxonomic order of each ASV. The outer two rings are heatmaps showing the log10-transformed average frequencies of each ASV in the native (outer ring) and barcoded (inner ring) 16S rRNA. Gray indicates not detected (N.D.) in amplicon sequencing data.

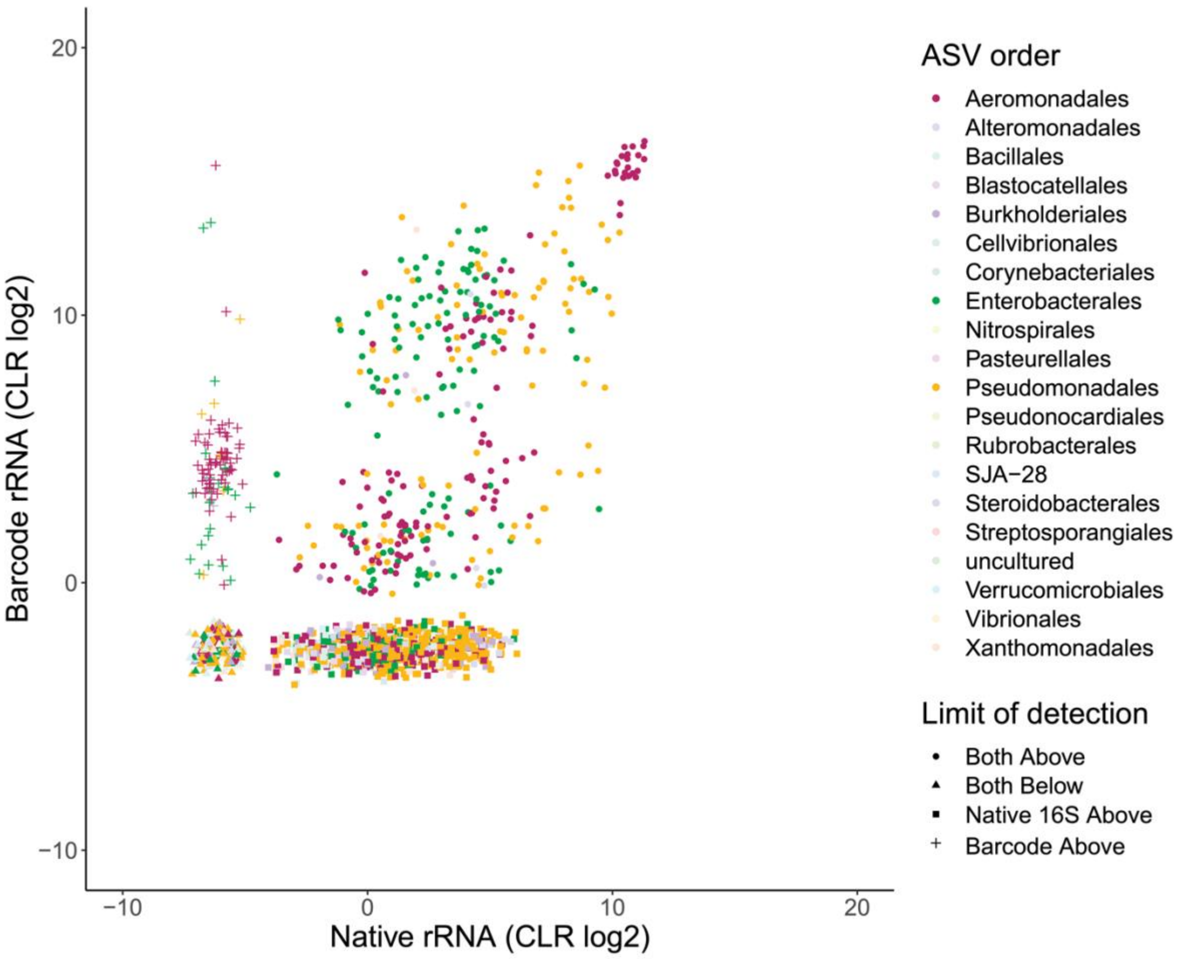


**Supplementary Figure 12. All ASVs detected from native rRNA and barcoded rRNA sequences.** Centered log-ratio (CLR) transformations were performed on barcoded and native 16S rRNA sequences from 6 replicate conjugations into a wastewater microbial community. Data coordinates represent the CLR values ($\log_2$) for each barcoded and native rRNA ASV from the same sample. Thus, each point represents an individual replicate of an observed ASV, with each ASV appearing six times on the scatter plot. Colors correspond to taxonomic orders, and shapes indicate which ASVs had: (circles) native and barcoded rRNA reads that were both above the limit of detection; (triangles) neither the native or barcoded rRNA reads above the limit of detection; (squares) only the native rRNA above the limit of detection; (plus) or only the barcoded rRNA above the limit of detection. These latter data represent ASVs that could not be detected via total rRNA sequencing, although they yielded a barcoded rRNA signal.

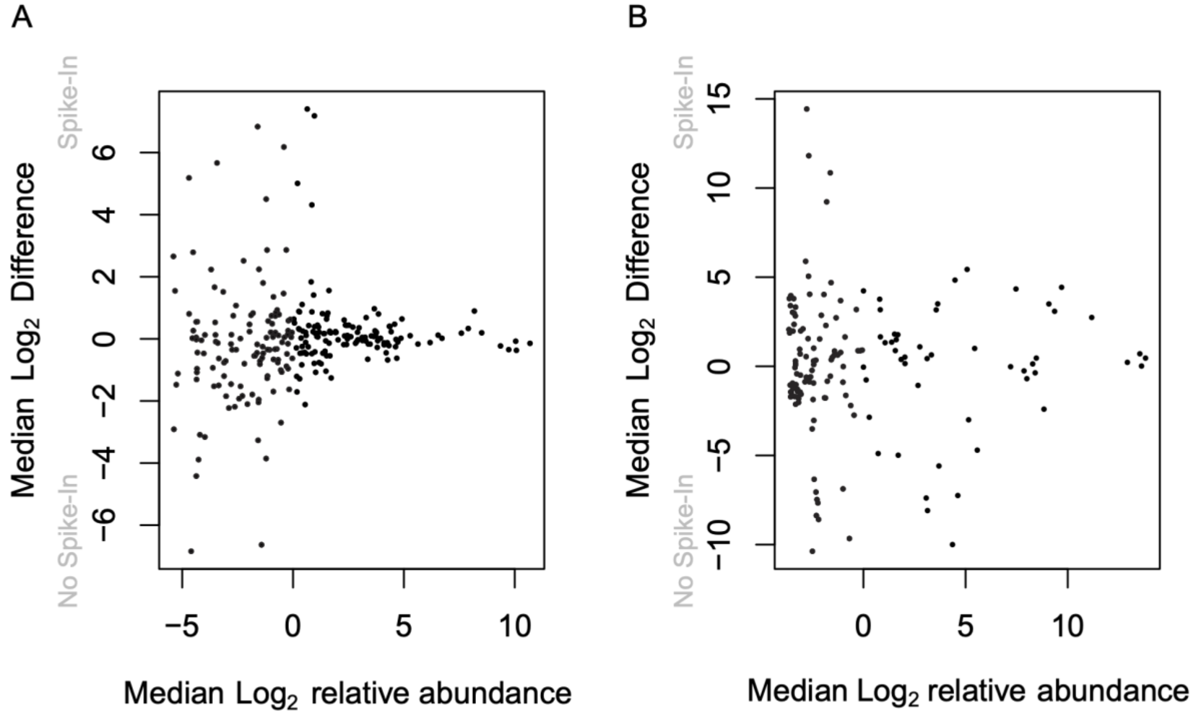


**Supplementary Figure 13. Barcoding in the wastewater community containing and lacking a non-native microbe spike in.** After conjugation, samples were split, and one was spiked with 400,000 *V. natriegens* to understand the intrinsic variance that exists in the wastewater population. Each sample, six in total, was analyzed for both ASV abundance (native rRNA) and DNA uptake (barcode rRNA) using amplicon sequencing. CLR transformations were performed for each sequencing run to compare results from different sequencing runs. The y-axis represents the difference between the median value spike and no spike in samples for each ASV, which was calculated by subtracting the *V. natriegens* spike in CLR median from the no spike in CLR median. The x-axis represents the median values of both the spike in and no spike in samples. Welch t-tests were performed comparing (**A**) native rRNA of spike in and no spike in samples, and (**B**) barcoded rRNA of spike in and no spike in samples**.** No statistically significant differences were observed between spike in and no spike in measurements with both comparisons.

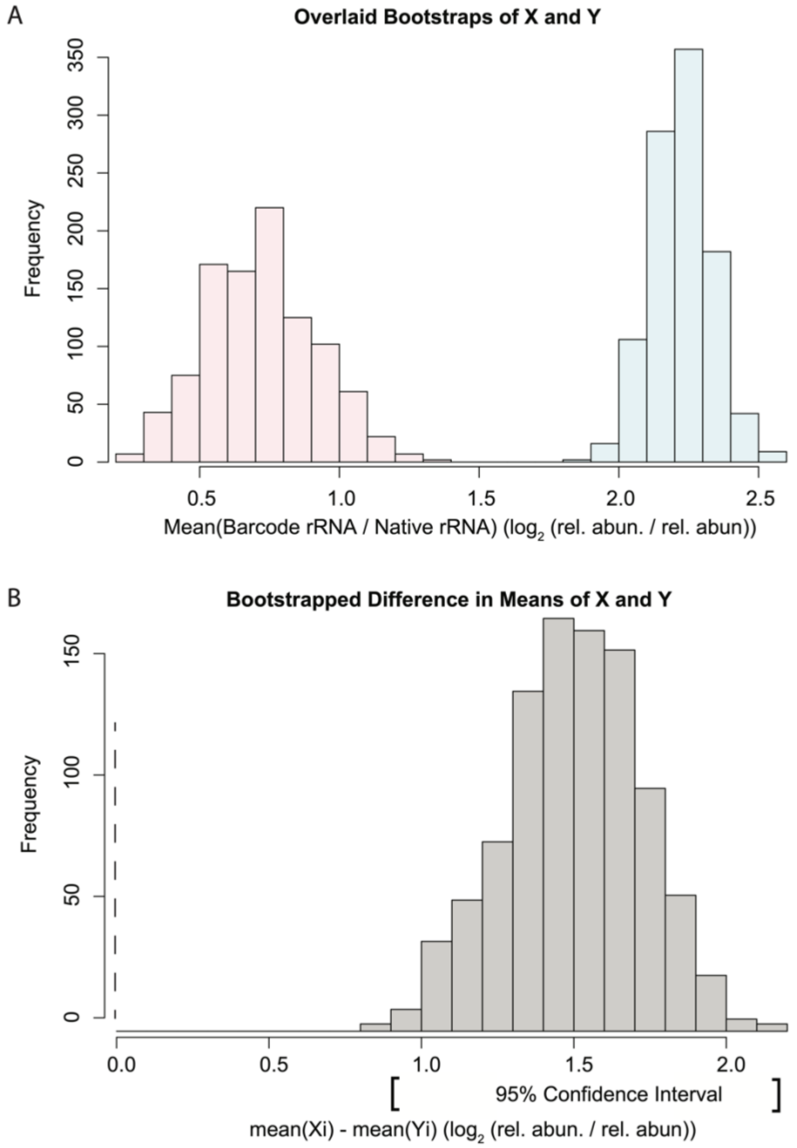


**Supplementary Figure 14. Bootstrapping reveals pairwise differences in ASV barcoding signals.** Provided is an example of a pairwise bootstrapped difference between means that was used to generate the data shown in Figure 3F. (**A**) The ratio of barcoded rRNA/native rRNA was computed by selecting from the six observed ratios with replacement, from which a mean ratio value is reported. For each ASV pair being compared, this process was repeated 1,000 times, creating the overlaid histograms in which 1,000 mean ratios are reported for each ASV. (**B**) A mean is randomly selected from the bootstrap pool from ASV *X* and from ASV *Y*, without replacement, and the difference between the two is calculated. The mean value of these 1,000 differences is what is reported for Figure 3F, and the 95% confidence interval is calculated and overlaid onto the graph. If the confidence interval does not cross over the dotted line, and therefore does not contain the value of 0 as a potential mean difference, the value was reported as certain and marked as such on Figure 3F. The above example shows that the difference between two ASVs is certain as it is outside the 95% confidence interval.

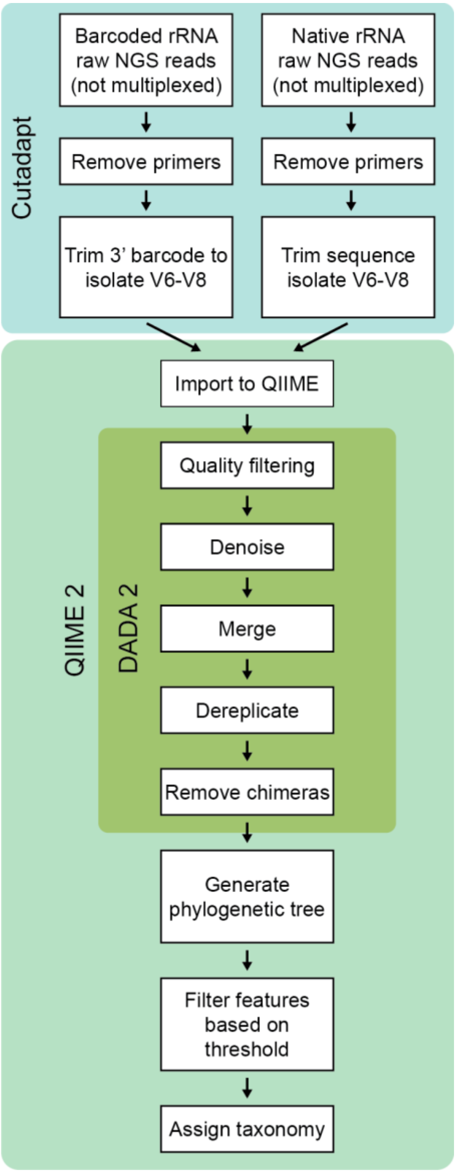


**Supplementary Figure 15. Workflow showing data processing to quantify barcoding.** Three computational tools were used for data processing: Cutadapt, QIIME 2, and DADA2. Cutadapt isolated sequences of identical homology (variable regions 6-8) from barcoded and native rRNA so that barcoded and native rRNA amplicons from the same organisms result in the same ASV. QIIME 2 with the DADA2 plugin was used to quality filter, denoise, demultiplex, and dereplicate reads, and also to remove chimeric sequences. QIME 2 was used for phylogenetic analysis.

### SUPPLEMENTAL TABLE LEGENDS

**Supplementary Table 1. Plasmids used in this study.** For the four categories of experiments, we indicate the plasmids constructed. For each plasmid, we note the use, the relevant data shown, and the plasmid architecture and features, including: (i) selectable marker, (ii) origin of replication, (iii) promoter, (iv) translation initiation sequences, (v) open reading frames, *e.g.*, sfGFP, and (vi) ribozyme details. CDF stands for CloDF13 origin of replication.

**Supplementary Table 2. Strains used in this study.** The genotypes for each microbe used are shown as well as their designations.

**Supplementary Table 3. Primers used for RT-qPCR and NGS.** The table indicates the primer-pairs used for PCRs along with an extra oligo (either probe for qPCR or reverse transcription primer for amplicon sequencing). The data corresponding to each specific primer pair is also noted for convenience. The nucleotides noted in lower case indicate partial Illumina adapter sequences for NGS sequencing. The AOS56B (1391R) primer serves as the universal primer for 16S amplification. AOS57B is the same as AOS56B used for reverse transcription but with the partial Illumina adapter added.

**Supplementary Table 4. Guide sequences used to target 16S rRNA.** The guide sequences that were attached to the ribozyme for each of the 16S splicing variants are shown. These sequences are complementary to 16S sequences, except with 1 mismatched ‘g’ (indicated in lower case) that is required by design, to make a ‘g:U’ wobble base pair with a ‘U’ in the 16S rRNA sequence.
