## Supplementary material for "Information storage across a microbial community using universal RNA memory": Methods

**Plasmid construction.** Table S1 lists all plasmids. Plasmids were constructed through either PCR cloning, Gibson assembly, or Golden Gate assembly (*1*, *2*). Sanger sequencing was used to verify all constructs.

**Strains, growth, and transformations.** Table S2 lists all strains. For all manipulations, single colonies were used to inoculate liquid cultures. *Escherichia coli:* cells were made competent by growing overnight in Lysogeny Broth (LB) medium at 37°C and 225 rpm, diluting 1:100 in fresh LB, and growing to an optical density (OD) of ~0.5. Cells were harvested by centrifugation, 3,000 g for ≥10 minutes (min), resuspended in TSS (1% w/v PEG3500, 0.5% V/V DMSO, 20 mM MgCl2 in LB) medium using a volume that concentrates 40 fold, and stored at -80°C in aliquots (200 µL). To transform, a cell aliquot was mixed with plasmid DNA (50 ng), incubated for 20 min on ice, heat shocked at 42°C for 45 seconds (s), grown for 1 hour (hr) in 2x YT medium (0.5 mL) at 37ºC and 225 rpm, and plated on LB-agar medium containing antibiotics. *Shewanella oneidensis MR-1:* cells were made competent by growing in LB medium for 17 hrs at 30°C and 250 rpm, were washed 3x by pelleting using centrifugation (7,900 g, 2 min) and resuspending in 10% glycerol (V/V), and ultimately resuspended in a volume of 10% glycerol that concentrates 15 fold. Unmethylated plasmid DNA (100 ng) purified from *dam⁻/dcm⁻ E. coli* (New England Biolabs) was mixed with freshly prepared cells (50 µL), incubated on ice for 1 min, and electroporated using 1.2 kV for ~5 milliseconds (ms) with a 0.1 cm cuvette. After recovery in LB medium (1 mL) for 2 hrs at 30°C and 250 rpm, cells were plated on LB-agar containing antibiotics. *Vibrio natriegens:* cells were made competent by growing in Brain Heart Infusion (BHI) medium with V2 salts (204 mM NaCl, 4.2 mM KCl, 23.1 mM $MgCl_2$) for 2 hrs 37°C and 250 rpm (*3*). When cells reached an OD of 0.5, they were pelleted, washed 3x times using electroporation buffer (680 mM sucrose and 7 mM $K_2HPO_4$, pH = 7), resuspended in a volume that concentrates 100 fold, and frozen in aliquots (50 µL) at -80°C. To transform, a cell aliquot was mixed with plasmid DNA (100 ng), incubated on ice for 1 min, and electroporated using a 0.9 kV pulse for ~5 ms using a 0.1 cm cuvette. After recovering in BHI medium (0.5 mL) supplemented with 680 mM sucrose and V2 salts for 1 hr at 37°C and 250 rpm, cells were plated on LB-agar containing V2 salts and antibiotics. *Pseudomonas putida:* *E. coli* MFDpir, a diaminopimelic acid (DAP) auxotroph (*4*), containing plasmid DNA was grown in LB supplemented with 0.3 mM DAP for 10 hrs at 37$^{o}$C and 250 rpm, while *P. putida* was grown in LB for 10 hrs at 37$^{o}$C and 250 rpm. After mixing these cultures at a 1:1 ratio, 50 µL was added to a nitrocellulose filter on top of LB-agar containing 0.3 mM DAP, and this plate was grown overnight at 25°C. The filter was placed in a 1.5 mL tube, and phosphate-buffered saline (PBS) was added (1 mL) and vortexed for 10 s. Cells were pelleted (6,800 g for 3 min), washed with PBS (1 mL), and plated on LB-agar containing antibiotics. *Pseudomonas stutzeri:* cells were made competent by growing for 16 hrs at 30°C and 250 rpm in LB. After harvesting cells by centrifugation (16,000 g, 2 min), they were washed 4 times by resuspending in 300 mM sucrose solution. Fresh cells (70 µL) that had been concentrated 86 fold were mixed with plasmid DNA (≥200 ng), incubated at 23°C for 1 min, and electroporated using a 2.5 kV pulse for ~5 ms with a 0.1 cm width cuvette. After allowing cells to recover in LB (1 mL) for 2 hrs at 30°C and 250 rpm, cells were plated on LB-agar containing antibiotics.

**Fluorescence measurements.** To assess ribozyme activity across diverse microbes, our visual reporter (pRAM18) was tested in *E. coli, S. oneidensis*, *P. putida,* and *V.*

*natriegens*. Single colonies containing pRAM18, 4 for each strain, were used to inoculate cultures containing antibiotics in a 2 mL 96-well block and grown for ~17 hrs at 1,000 rpm in a VorTemp 56 (Labnet) shaker. LB medium was used for all cultures, except *V. natriegens* which used LB containing v2 salts. After inoculating fresh LB (1:50 dilution), cultures were grown for 3 to 8 hrs at 30 or 37°C and 1,000 rpm. Whole-cell fluorescence was performed using cells (50 µL) mixed with PBS (50 µL) in a 96-well plate. Optical density (OD) at 600 nm and sfGFP fluorescence (FL) were measured using a Tecan Spark (excitation: 485 nm; emission: 510 nm). OD and FL values were corrected by subtracting the corresponding mean values of the media blank. The FL to OD ratio (FL/OD) was calculated for each well, which was derived from a single colony. Normalized FL/OD was calculated for each colony by dividing the mean of a specified condition (described in each figure legend) and each data point presented. For analysis of conjugative vector repression, we transformed a vector that expresses mScarlet under control of the CymR promoter (pRAM24) into *E. coli* MG1655 containing or lacking a CymR expression plasmid (pRAM25). Single colonies were used to inoculate LB cultures (1 mL) containing antibiotics. After growing for 18 hrs at 37°C and 650 rpm, OD and FL were measured (excitation: 569 nm; emission: 610 nm). For flow cytometry analysis, overnight cultures of cells harboring pRAM18 and appropriate controls (pRAM16, pRAM17) were diluted 1:100 and analyzed using a Sony Biotechnology SH800 Cell Sorter. In total, 50,000 events were collected per sample (excitation: 487 nm; emission: 525 nm). Flowjo (BD Biosciences) was used to process flow cytometry data. An elliptical gate was used to isolate the densest population on the plot containing SSC-H vs FSC-H channels, containing ~80% of the population. A second gate was used on the FSC-H vs FSC-A for doublet discrimination.

**Growth measurements.** Single colonies (n = 12 each) of *E. coli* MG1655 harboring plasmids that express the visual reporter (pRAM18), a sfGFP positive control (pRAM17), and an empty vector (pRAM16) were used to inoculate LB cultures (1 mL) containing antibiotics. After growing cultures overnight at 30°C and 650 rpm, 1 µL was used to inoculate LB medium (99 µL) containing antibiotics in Nunc Edge 2.0 96-well plates (Thermo Fisher). Cells were grown for 24 hrs at 30°C with double orbital rotation (3 mm amplitude) in a Tecan M1000 while measuring the OD every 10 min. Maximum specific growth rates were obtained using Growth Curve Modeler (*5*).

**RNA extraction.** *E. coli, S. oneidensis*, *P. putida, P. stutzeri,* and *V. natriegens* transformed with plasmids for expressing different cat-RNA were grown in LB as described for sfGFP fluorescence. Stationary phase cultures were diluted 1:100 into LB medium (1 mL) containing antibiotic, except *V. natriegens* which was diluted into LB medium containing V2 salts. After growing for ~4 hrs, cells were pelleted (4,000 g, 5 min). The Maxwell® Purefood GMO and Authentication Kit (Promega) was used to extract total DNA and RNA using an automated magnetic bead purification with a Maxwell® RSC 48 instrument. In brief, cell pellets were resuspended by vortexing in CTAB buffer (1 mL), heated for 5 min at 95°C, incubated at 23°C for 2 min, and vortexed. Proteinase K (40 µL, 20 mg/mL) was added, vortexed, and incubated at 70 °C for 10 min. Lysed samples (300 µL) were added to the receiver well of the extraction cartridges along with lysis buffer (300 µL), and elution buffer (50 µL) was used to obtain total RNA and DNA. DNA was then removed by adding 2 µL of Turbo DNAse (Thermofisher) per 10 µg RNA in a 50 µL reaction volume. The sample was cleaned up using a Turbo DNA free kit (Invitrogen) by adding the chelating bead suspension (5 µL), incubating for 10 min while agitating every 2 min to resuspend the beads, and centrifuging at 10,000 g for 3 min. To obtain the RNA concentrations, the resulting supernatant (47 µL) was analyzed using a Qubit™ Broad Range (BR) Assay Kit. To ensure that all inhibitors from the kit were diluted, all samples were adjusted to an RNA concentration that was 10 fold lower than the sample having the lowest concentration following purification.

**Reverse transcription quantitative PCR (RT-qPCR)**. Primers (Table S3) and probes used for RT-qPCR were designed using Primer3plus with user-defined thermodynamic parameters (*6*). RT-qPCR reactions quantifying barcoding by different cat-RNA designs were performed using a dye-based qPCRbio Sygreen 1-step kit (PCR Biosystems, PB25.11) in a Quantstudio-3 qPCR thermocycler (ThermoFisher). RNA template (4 µL) was used in a 10 µL total reaction volume with 0.4 µM of each primer and the provided RTase Go reverse transcriptase (0.5 µL). To determine an initial concentration, RT-qPCR amplification curves were analyzed using LinRegPCR (*7*), which background subtracts, log-transforms, and finds the linear phase of the amplification curve. To compare the concentration of native and barcoded 16S rRNA across microbes and measure the stability of barcoded rRNA, RT-qPCR reactions were performed with probes using PCRbio Probe 1-step Detect master mix (PCR Biosystems, PB25.41). RNA template (4 µL) was added into a 10 µL total reaction volume with 0.4 µM of each primer and 0.2 µM of each probe along with RTase Go reverse transcriptase (0.1 µL). For the same samples, the 16S rRNA quantification was performed in a 10 µL reaction using a dye-based qPCRbio Sygreen blue mix (PCR Biosystems, PB20.15) with RTase Go reverse transcriptase spiked in (0.5 µL). A custom R script was used to convert the Cq data exported from Thermo Fisher Quantstudio software into absolute quantities in copies/µL template using standard curves. In all cases, sample concentrations were converted to copies/µL template using standard curves made with known concentrations of commercially-synthesized DNA (IDT) or purified PCR products. Scripts are available on GitHub (github.com/ppreshant/qPCR-analysis).

**RNA stability measurements.** To measure RNA stability, *E. coli* harboring vectors that express GFP (pRAM17) or the cat-RNA that targets U1376 (pRAM23) were grown to stationary phase in LB medium (3 mL) containing antibiotics, diluted 1 to 100 into fresh LB (3 mL), and grown for 3 hrs at 37°C. Cells were diluted to an OD of 0.5 into fresh LB medium (5 mL) containing or lacking the transcription inhibitor rifampicin (150 ng/µL). Samples (750 uL) harvested from these cultures after 0, 10, 30, 60, and 180 min incubations were frozen prior to RNA extraction and RT-qPCR analysis.

**Universal cat-RNA guide design.** To identify possible cat-RNA designs, a 16S rRNA sequence alignment was created using four *Gammaproteobacteria* strains and one *Bacillus* strain, including *E. coli* K-12 MG1655 (NCBI: NC_000913.3, bases 223771-225312), *P. stutzeri* NCTC10475, (NCBI: LR134482.1, bases 939610-941134), *S. oneidensis* MR-1, (NCBI: AE014299.2, bases 46116-47649), *V. natriegens* NBRC 15636, (NCBI: CP009977.1, bases 55-1617), and *Bacillus subtilis subsp. subtilis* 168, NCBI: CP053102.1, bases 30279-31832). First, we identified the uracils that were conserved across all of the 16S rRNA from each species. Those uracils within 35 nucleotides (nt) of the 3' end of the 16S rRNA were discarded because we set a minimum sequence length of 35 nt for complementary sequence to the 50 nt guide RNA. All uracils within 5 nt of the

5’ end of the 16S rRNA were also discarded, because they could not be sequenced if barcoded. Second, we used Clustal Omega multiple sequence alignment to establish the conservation in the 5 nt upstream to the conserved uracils (*8*), where the IGS in the cat-RNA will be designed to anneal, which is designated the P1 stem. To maximize IGS binding in synthetic cat-RNA designs, only those uracils and adjacent P1 stems with 100% sequence conservation were used for further analysis. Third, we created a consensus sequence corresponding to the 50 nt downstream of every completely conserved P1-uracil, which corresponds to the region that is complementary to the synthetic guide sequences in cat-RNA. To do this, we used BioPython’s “dumb_consensus” with a threshold of 0.70, and denoted any base pairs as ambiguous *N* if below this threshold (*9*). The consensus score $S(x)$, which ranged from 0 to 1, was calculated using the consensus sequence as follows:

$$S(x) = \frac{\sum_{i=1}^{i=n} \alpha(x_i)}{n}$$

where $x_i$ is the identity of the base at location $i$ of the consensus sequence having length $n$, and $\alpha(x_i)$ is 1 if $x_i$ = A, U, C, or G and $\alpha(x_i)$ is 0 if $x_i$ is N. As such, the consensus score provides a simple metric of sequence conservation. Finally, we designed complementary cat-RNA guide sequences (50 nt) based on 4 sequences with the high consensus scores, which were used to guide the construction of plasmids that express four different cat-RNA. These are named based on the uracil targeted for the trans splicing reaction in *E. coli* 16S rRNA, which included U17, U891, U1376, and U1490 (Supplementary Table 4). Community conjugation studies all used cat-RNA U1376. The variable regions in *E. coli* 16S rRNA used for comparisons were previously reported (*10*).

**Amplicon sequencing of barcoded rRNA in *E. coli*.** To convert extracted RNA to DNA for amplicon sequencing, purified RNA (~500 ng) was combined with primers (0.5 µL each, 2 µM), dNTPs (0.5 µL, 10 mM), and water to a final volume of 6.5 µL. RNA was denatured at 65°C (5 min), cooled on ice for 5 min, and the following reagents (Thermofisher) were added: dithiothreitol (0.5 µL, 100 mM), 5x first strand buffer (2 µL), RNAseOUT ribonuclease inhibitor (0.5 µL, 40 U/µl), Superscript III reverse transcriptase (0.25 µL, 200 U/µL), and water (2.5 µL). Reverse transcription was carried out at 55°C for 1 hr, heat inactivation was done at 75°C for 15 min, and the resulting cDNA was stored at -80°C. PCR amplification was performed in a 25 µL reaction by combining cDNA (0.5 to 1 µL), forward and reverse primers (0.125 µL each, 100 mM), dNTPs (0.5 µL, 10 mM), 5x Q5 reaction buffer (5 µL), and Q5 DNA polymerase (0.25 µL, 2 U/µL). The following PCR protocol was used: 1 cycle of 98°C for 1 min, 25 cycles of 98 °C for 10 s, 60°C for 25 s, and 72°C for 20 s, and 1 cycle of 72°C for 2 min. An aliquot of the product (3 µL) was analyzed on a 1% TAE-agarose gel to visualize the amplicon. Gel analysis confirmed the presence of the expected barcoded product (552 bp) along with an additional side product. All potential PCR products having a range of sizes were purified using a silica spin column (Epoch) and sequenced by Genewiz using their Amplicon-EZ protocol.

**Barcoding specificity analysis.** Deep sequencing of barcoded rRNA in *E. coli* was returned as paired-end reads, with the forward and reverse reads in separate files. Even though all PCR products are expected to contain a splice site, since they were generated using one primer that anneals to the 16S rRNA and a second primer that anneals to the barcode, the forward and reverse sequencing reads will only detect the splice site in cases where it is proximal to the sequencing adapters, since Amplicon-EZ sequencing yields short reads. The reverse reads are all expected to begin with the barcode sequence, which is proximal to the splice site junction between 16S rRNA and the barcode (<79 bp), so all of these reads are predicted to contain splicing information. The forward reads begin with 16S rRNA sequence information, so only a subset of these reads are expected to yield splicing information, given their wide range of potential sizes. For example, forward reads of small PCR products (<250 bp) are expected to contain a splice site, while reads of longer PCR products (>250 bp) are expected to yield sequences that are completely 16S rRNA sequence as the reads do not extend to the splicing site. Due to these constraints, only the reverse sequencing reads were used for the identification of the splice site within 16S rRNA. Cutadapt was used to first find reads with the reverse primer sequence (AACCTTCGGGCATGG) used for PCR, and then it was used to remove the primer sequences from these reads using the default parameters (*11*), with the exception of setting the minimum overlap to 5 and minimum length to 1. The forward PCR primer site (AACGCGAAGAACCTTAC) was also removed for small PCR products where the reverse read covered the entire barcode, 16S rRNA, and forward PCR primer sequence. If no reverse primer sequence was identified in a read, the read was discarded. Using Cutadapt, the barcode sequence was identified and removed in each read to yield each 16S rRNA sequence ending in the corresponding splice site. If no barcode sequence was identified in a read, the read was discarded. Using a custom python script, identical reads were then dereplicated and the reverse complement of each unique read was generated. The splice site was determined by aligning each unique read to the *E. coli* 16S rRNA sequence. The point where the alignment ended was designated the splice site. The P1 stem was defined as the nucleotides immediately upstream of the splice site in the 16S rRNA sequence.

**Wastewater sampling.** Mixed liquor (50 mL) was collected on March 8th, 2022 from the aeration basin of the West University Wastewater Treatment plant in Houston, Texas. The sample was immediately placed on ice and transported to the laboratory where conjugation was performed within 4 hrs of sampling. Prior to conjugation, the wastewater sample was homogenized by vortexing it. All experiments that involved wastewater samples were performed at room temperature.

**Wastewater conjugations.** *E. coli* MFDpir was used as a donor strain for conjugation (*4*). Donor cells transformed with vectors that express CymR (pRAM25) and the cat-RNA designed to target U1376 (pRAM24) were grown to late log phase (10 hrs) in LB (3 mL) containing 0.3 mM DAP and antibiotics. These cells (OD = 1) were washed 3x times with PBS (1 mL) to remove antibiotics and mixed with the wastewater microbial community at a 1:1 (V/V) ratio. An aliquot of each mixture (50 µL) was added onto a nitrocellulose filter that had been placed on LB-agar medium containing DAP (0.3 mM). After incubating at 25°C for 24 hrs, filters were removed, placed in a 1.5 mL tube, and washed with PBS (1 mL). Cells were recovered by vortexing (10 s) and centrifuging (3 min, 6800 g). The filter paper was removed, and the cells were washed 2x using PBS (1 mL). For RNA extraction, a modified protocol of the Maxwell® RSC PureFood GMO and Authentication Kit (Promega) was used to extract nucleic acids from the cell pellet. Alterations included excluding RNAse, incubating the proteinase K at 37°C, and using 50 uL of elution buffer per sample. Samples were DNAse treated prior to reverse transcription of RNA to cDNA, as described above in “RNA Extraction.”

**Amplicon sequencing of the wastewater microbial community native and barcoded 16S rRNA**. Total RNA was converted to cDNA as described for *E. coli*. These cDNA libraries were PCR amplified in 100 µL reactions by combining cDNA (1 µL), forward and reverse primers (0.5 µL each, 100 µM), dNTPs (2 µL, 10 mM), 5x Q5 reaction buffer (20 µL), and Q5 DNA polymerase (1 µL, 2 units/µL). After dividing samples into four aliquots, touchdown PCR was used: (step 1) 98°C for 2 min followed by 12 cycles of 98°C for 30s, 72°C for 45 s with a 1°C/cycle decrease in temperature, and 72°C for 1 min; (step 2) 2 cycles of 98°C for 30s, 60°C for 45 s, and 72°C for 1 min; (step 3) 19 cycles of 98°C for 30s and 72°C for 1 min 45 s, and (step 4) 72°C for 2 min and 4°C for 15 min. PCR products were separated on a 1% agarose gel, the expected products were gel purified (516 bp for barcoded rRNA and 505 bp for native rRNA), and these products were sequenced using the Amplicon-EZ service (Genewiz). When performing agarose gel analysis of the PCR product, unexpected bands were observed, which were removed by gel purification. All reactions yielded >224,000 reads.

**Data analysis pipeline.** The workflow for data analysis is shown in Supplementary Figure 15. Cutadapt was used to trim the raw sequencing reads so that both native and barcoded rRNA sequences comprised the same homologous regions (*11*). To do this, the forward primer sequences (AACGCGAAGAACCTTAC) for both barcoded and native rRNA amplicons were removed, as well as the reverse primer sequences, which differed for the native (TGACGGGCGGTGWGTRCA) and barcoded (GTTTCATGTGATCCGGATAAC) barcoded amplicons. For the barcoded rRNA amplicon reads, the barcode sequence (ATGGTGTTCAATGCTTTTCCC) was removed. Because the native 16S rRNA amplicon contained a short sequence (ACGTTCCCGGGCCT) after the splice site, this sequence was also trimmed to make the two amplicons completely homologous. Reads were required to have a minimum amount of the trimmed sequence align to the end of the read (5 nt for the primers and the sequence after the splice site, 10 nt for the barcode) and could not have mismatches that exceeded a threshold (10% for primers and barcoded, 14% for the sequence after the splice site). Any reads that lacked any of these trimmed sequences or exceeded the error tolerance were discarded. After trimming, NGS reads were imported into QIIME 2 version 2021.11 (*12*). Sequence quality assessment using `demux summarize` revealed that sequences were high quality and required no additional end trimming. The DADA2 plugin was used to quality filter, denoise, demultiplex, and dereplicate the reads. Chimeras were also removed using DADA2 (*13*). This processing resulted in a feature table showing the sequences of each unique amplicon sequencing variant (ASV), their relative abundances, and the total number of ASV counts observed in each sample. A phylogenetic tree of all ASVs was generated with a fragment-insertion approach using the plugin `fragment-insertion sepp` (*14*). Both the tree and the feature table were filtered to only contain ASVs that appeared in at least three reads across at least two samples. This limit of detection was required for all analysis. Finally, taxonomy was assigned to each ASV using `feature-classifier classify-sklearn` with a pre-trained classifier (*15*–*17*).

**Comparing native and barcoded rRNA levels across ASVs.** Sequence read abundances from 12 sequencing datasets generated from 6 replicate samples (*i.e.*, paired native and barcoded rRNA for each replicate) were vectorized and linearly transformed with centered log-ratio (CLR) transformations using ALDEx2 software (*18*). The log transformation is base 2, reporting fold changes. To understand the variance within our consortium conjugation data, samples were split into two pools, and ~400,000 *V. natriegens* cells were added directly to one sample prior to RNA extraction, which is not found in the wastewater consortium. These paired samples were compared using the ALDEx2 built in function difference between (diff.btw). From there, custom software was used to further analyze the CLR transformed vectors. Sequencing runs were paired, with each replicate tested for native and barcoded 16S rRNA. To determine the relationship between rRNA barcoding and native rRNA levels for each ASV, all samples that yielded barcoding and native reads above the limit of detection were fit to a linear model. For some ASVs, native rRNA was detected but not barcoded rRNA, while other ASVs yielded barcoded rRNA signals but not native rRNA signals (Supplementary Figure 12).

**Determining confidence intervals for barcoding efficiencies.** To quantify the relative barcoding efficiency for each ASV (*i.e.*, the barcoded to native rRNA CLR ratios), only those ASVs which yielded detectable barcoded and native rRNA in all six replicates were further analyzed. This represented ASVs from three of the orders observed in the wastewater microbial community. To compare barcoding between ASVs, a bootstrap approach (n = 1,000) was used to compute the difference between means for each pairwise comparison, *i.e.*, 28 total ASVs, and the magnitude of each difference is reported as a heatmap. To assess the certainty that a given ASV pair has distinct barcoding levels (Supplementary Figure 14), the 95% confidence interval was calculated using the bootstrap analysis. In cases where this 95% confidence level does not include a value of 0 as a potential mean difference, the pairwise difference of the ASVs was reported as certain within the heatmap.

**Statistical analysis of wastewater microbial community data.** A Welch t test was used to compare samples containing or lacking a *V. natriegens* spike in using the ALDEx2 software (*18*), with a p value < 0.1 being used as a threshold for significance. All other p-values were determined by performing two-tailed, independent t tests with using p < 0.05 as a threshold for significance.

**Calculation of free energies for cat-RNA and target RNA binding.** A locally-installed version of the Nucleic Acids Package (NUPACK) version 4.0.0.27 was used to predict free energy terms using default parameters (*19*).

**Shannon entropy calculations.** To calculate the Shannon entropy $H(x_i)$, we first aligned *m* 16S rRNA sequences to *E. coli* 16S rRNA. At each position *i* in each 16S rRNA sequence *j*, we calculated $P_c(x_i)$ as the sum of the normalized probabilities $p_c(x_{i,j})$ of nucleotide *c* (*i.e.*, A, C, G, U, or gap) in each sequence *j*, where there are *n* = 5 possible states considered. $P_c(x_i)$ was calculated using *m* sequences as:

$$P_c(x_i) = \frac{\sum_{j=0}^{m-1} p_c(x_{i,j})}{n}$$

For sequences with a single canonical bp (A, C, G, U, or gap) at position *i*, the probability $p_c(x_{i,j})$ of finding residue *c* in a given sequence was designated 1. However, if positions presented ambiguity, *e.g.*, a purine (R), then we designated each base pair (A, G) as having an equivalent probability of 0.5. $H(x_i)$ was calculated at each position *i* as:

$$H(x_i) = \sum_{c \in [A,U,G,C,-]} -P_c(x_i) * log_2(P_c(x_i))$$
